# To remain modern, the coexistence program requires modern statistical rigor

**DOI:** 10.1101/2022.12.28.522056

**Authors:** David W. Armitage

## Abstract

A recent study by Van Dyke et al.^1^ paired experimental drought manipulations with demographic models and trait measurements to project major shifts in coexistence among a number of annual plant taxa. However, re-analysis of the data under alternative, more predictive competition models reveals that the authors’ original conclusions are very sensitive to slight variations in model form. Furthermore, propagating model parameter error into coexistence predictions results in relatively weak support for the majority of coexistence shifts predicted by the authors’ original model. These results highlight the need for increased statistical rigor when treating binary predictions of species coexistence as observed experimental outcomes, as is commonly practiced in empirical coexistence studies.

## Main Text

Forecasting the effects of precipitation change on plant communities is a paramount challenge. The theoretical framework of *modern coexistence theory* (MCT) has been used to predict the joint contributions of niche and fitness differences (ND and FD, respectively) to competitive outcomes and species coexistence under future precipitation projections^1, 2^. By tracking the demographic rates of plants in experimental communities receiving either reduced or ambient precipitation over one growing season, Van Dyke *et al.^1^* use this framework to show that moderate decreases in water availability will shift the predicted coexistence outcomes of 10 of the 15 annual plant species under study, and that these shifts are more likely in functionally diverse communities. Such a finding is noteworthy as functional diversity is anticipated to contribute to the maintenance of ecosystem services and is therefore often a desired outcome of restoration and conservation projects^3^.

However, Van Dyke *et al.’s* results depend heavily on the key assumption that species pairs which satisfy the inequality *ρ* < *k_j_*/*k_i_* < 1/*ρ* (where *ρ* denotes niche overlap and *k_j_*/*k_i_* fitness differences) will stably coexist^4^. In the absence of independent data to benchmark the empirical accuracy of this inequality, it is imperative that the estimates of ND and FD be statistically robust. To this end, Van Dyke et. al^1^ omit some important statistical analyses such as model selection and error propagation, which affect their results concerning drought-mediated shifts in coexistence outcomes and relationships between trait and fitness differences.

The first issue is that of model specification. There are many ways to write phenomenological competition models which are nearly equivalent in both assumptions and complexity, but that assume slightly different functional forms of density-dependence^5^. Following the authors’ prior work^6, 7^, Van Dyke *et al.* assume that a simple form of the Beverton-Holt (BH) competition model best describes the dynamics of their system. Given that the output of the analysis is a theoretically-motivated prediction (coexistence or competitive exclusion), and there is no *a priori* basis to strongly favor the BH model over similar alternatives, then it follows that the model with the best predictive accuracy on withheld data should be the one most trusted to generate parameter estimates used in subsequent predictions and analyses.

To investigate the sensitivity of model choice on the results, I used a Bayesian approach to sample the posterior distributions of competition (*α_ij_*), growth rate (*λ_i_*), and treatment effect parameters for seven different alternative competition models of similar complexity. For each focal species, and using weakly informative priors with the same constraints as those used by the authors, I ran eight Markov chains of length 10000, discarding the first 50% as warmup samples. After confirming MCMC convergence and that posteriors and resulting ND and FD estimates of the BH model matched those from Van Dyke *et al.,* I fit six additional model forms to the same data^5, 6, 8^. Comparing models using the Watanabe-Akaike Information Criterion (WAIC) — a complexity-penalised measure of a model’s out-of-sample predictive performance^9^ — I identified three models that better predicted independent data than the BH model of Van Dyke *et al.* (Table 1). A slightly modified BH model (no. 7) offered the best improvement of predictive ability and stability compared to other high-ranking models.

**Table 1.** Comparison of various competition models of density-dependent fecundity, *F_i_* using WAIC. Values shown are mean ± standard deviation across species. Here, model no. 7 has the best predictive ability and is a significant improvement over the BH model (no. 5). Alternative model comparisons using standard AIC_c_ and BIC on maximum-likelihood fits return quantitatively consistent results.

| Model | $\langle \text{WAIC} \rangle$ | $\langle \text{SE.WAIC} \rangle$ | $\langle \Delta \text{WAIC} \rangle$ |
| --- | --- | --- | --- |
| (1) $F_i = \lambda_i$ | $488 \pm 117$ | $16.6 \pm 4.9$ | $108.0 \pm 63.8$ |
| (2) $F_i = \lambda_i - \alpha_{ii}N_i - \alpha_{ij}N_j$ | $423 \pm 81$ | $19.3 \pm 3.7$ | $42.6 \pm 21.1$ |
| (3) $F_i = \lambda_i e^{-\alpha_{ii}N_i - \alpha_{ij}N_j}$ | $423 \pm 81$ | $19.3 \pm 3.7$ | $42.8 \pm 21.1$ |
| (4) $F_i = \lambda_i e^{-\alpha_{ii} \log(N_i+1) - \alpha_{ij} \log(N_j+1)}$ | $383 \pm 69$ | $21.5 \pm 5.3$ | $2.7 \pm 2.5$ |
| (5) $F_i = \lambda_i / (1 + \alpha_{ii}N_i + \alpha_{ij}N_j)$ (BH Model) | $392 \pm 68$ | $21.0 \pm 5.4$ | $11.6 \pm 10.1$ |
| (6) $F_i = \lambda_i / (1 + N_i^{\alpha_{ii}} + N_j^{\alpha_{ij}})$ | $382 \pm 70$ | $21.5 \pm 5.2$ | $1.7 \pm 1.8$ |
| (7) $F_i = \lambda_i / (1 + \alpha_{ii}N_i + \alpha_{ij}N_j)^\theta$ | **$381 \pm 70$** | **$21.8 \pm 5.4$** | **$1.2 \pm 2.1$** |

Using 1000 posterior draws of *λ_i_*, *α_ii_*, and *α_ij_* from model 7, I calculated ND and FD for each species pair and assessed whether these draws satisfied the aforementioned coexistence inequality (Figure 1). Performed over the set of posterior draws for each species pair, this process generates a distribution of coexistence probabilities conditioned on the model, priors, and data. I then calculated the probability that a switch in coexistence outcomes had occurred between treatments. This probability, *p*(switch), is defined as *p*(*C_i_* ∩ *E_j_*), *i* ≠ *j* where *p*(*C_i_*) is the probability of coexistence in the precipitation treatment *i* with the highest coexistence probability, and *p*(*E_j_*) is the probability of exclusion (= 1 – *p*(*C_j_*)) of the other treatment, *j*. Two key findings emerged. First, that coexistence predictions for most species pairs are model-dependent and therefore sensitive to subjective choices made during model selection. Second, of the original 10 species pairs predicted to have switched coexistence outcomes between treatments, only three such switches are now predicted at probabilities greater than 0.5 (Figure 1). Model 4, which also had a better fit to the data, only predicted one species pair (PL-FE) to switch outcomes. However, no additional species pairs beyond those already identified by the authors were predicted to have switched under these models.

**Figure 1.**
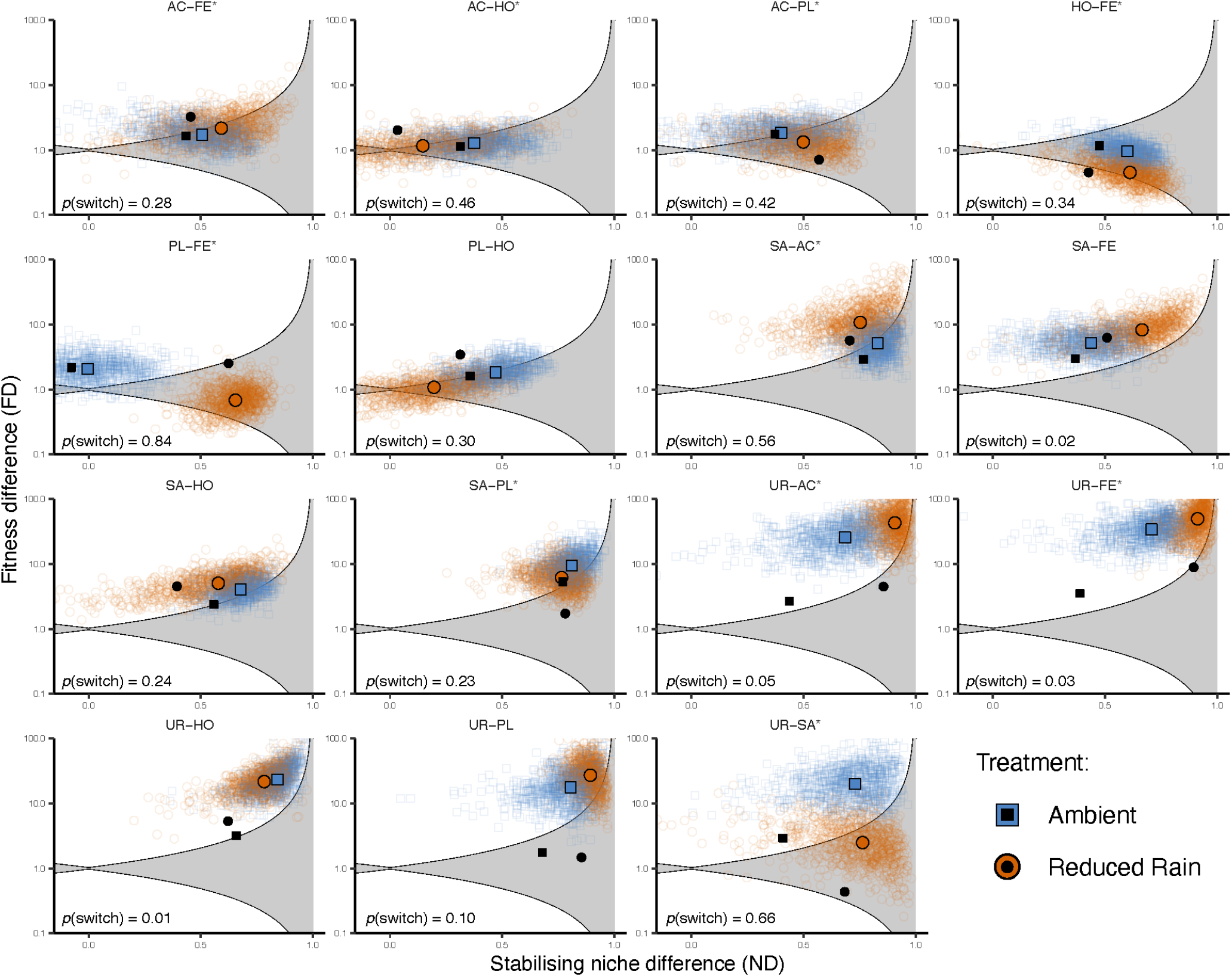
Predictions of coexistence outcomes for 15 species pairs. Points falling inside of the shaded region are those that satisfy the coexistence inequality. Colored points represent posterior draws from model 7. Solid colored shapes are median posterior estimates from this model and black shapes are authors’ eestimates from the BH model. For each panel, the probability that a switch between coexistence and exclusion having occurred is also shown. Asterisks denote species pairs predicted to have experienced coexistence shifts in the original analysis.

Carrying the median posterior estimates of model 7’s niche and fitness differences forward through the remaining analyses results in the loss of statistically significant differences between competition and demographic differences between treatments (Figure 2a). Further, and perhaps most importantly, changes in FD between treatments are no longer significantly positively associated with the functional trait differences between species pairs (Figure 2b). We are left to conclude that under a competition model with better fit to the data than the standard BH (e.g., models 7 and 4 in table 1), most of the major conclusions concerning drought-mediated shifts in coexistence disappear.

**Figure 2.**
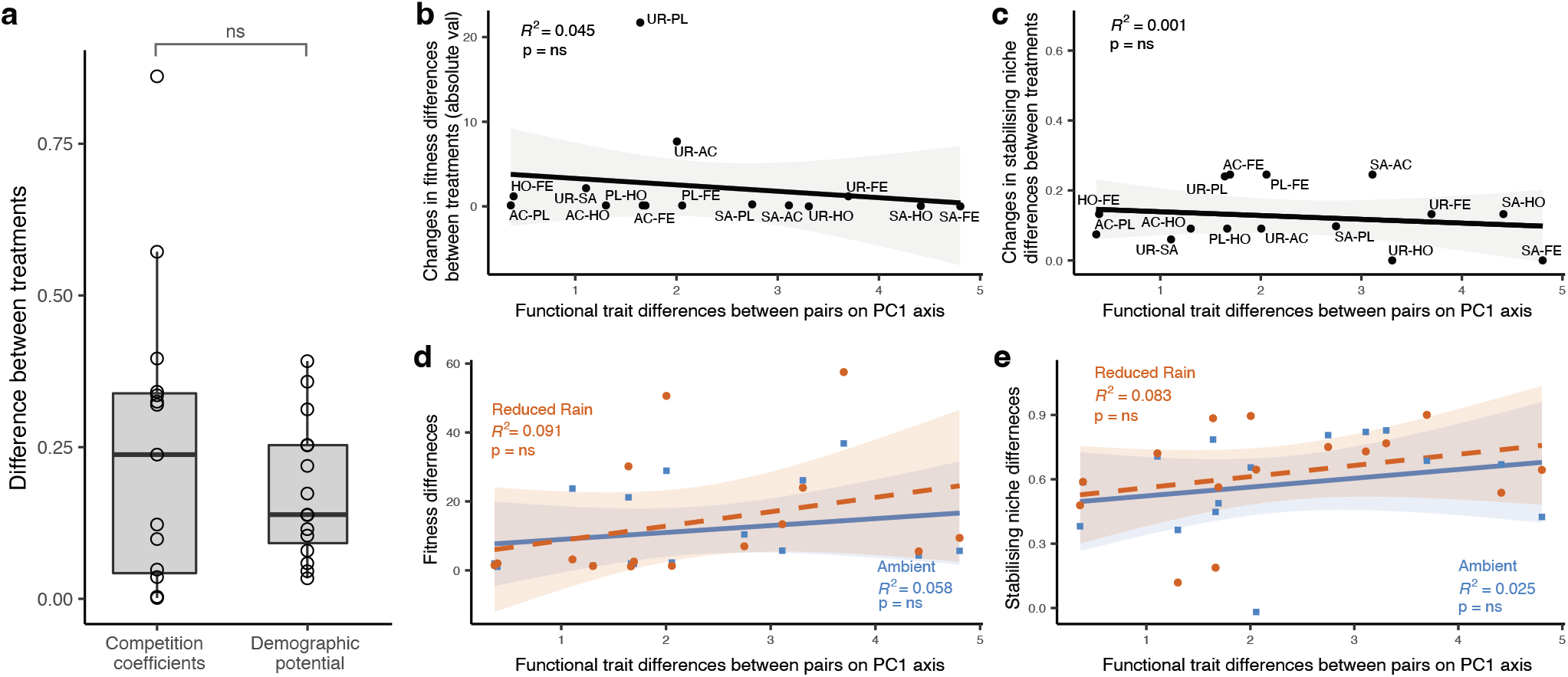
Using the coefficients of model 7 has the effect of removing statistically significant trends in a) the relative magnitudes of treatment differences in demographic potential and competition coefficients (*t* = −1.02, *p* = 0.32) and, b-e) the positive relationships between functional trait distances and treatment-driven distances in FD. ND results remain unchanged.

It could be argued that despite support for better-performing models, the standard BH model has been so widely used, that it should be the considered the preferred model to standardise comparisons across studies. However, even in the rare cases where these studies present error estimates for ND and FD, decisions concerning coexistence outcomes are rarely made with the same level of statistical rigor. The statistical philosophy employed throughout the analyses of Van Dyke *et al.* is inconsistent. Most of the authors’ analyses present statistical evidence in the form of null hypothesis tests with a type I error tolerance of 5%. However, this is abandoned in one key area — decisions about whether or not a species pair is scored as coexisting. Instead, the authors use median values of ND and FD from a non-parametric bootstrap to make binary predictions with the error tolerance of 50%. While error bars are kindly provided in a supplemental figure, many clearly transect the coexistence boundary defined by ND and FD and therefore do not enjoy the same level of statistical support as their other conclusions. Though there are no agreed-upon methods for what a null hypothesis test of coexistence predictions should entail, I suggest that propagating error either through the non-parametric bootstrap samples or posterior draws of parameters can quantify support for these predictions.

I illustrate this by using the BH model’s posterior draws to propagate error through to ND and FD estimates. Median values of these draws closely matched the authors’ maximum likelihood estimates (Figure E1). I then calculated the probability of coexistence for each species pair × treatment combination and, subsequently, the probability that a switch between coexistence and exclusion has occurred. For the species pairs identified as having switched between the two outcomes, the average probability of this switch was 0.61 ± 0.11, and the joint probability that all of these pairs had switched was < 0.01. In no case was the probability of switching greater than 0.9, which is what would be expected if, under a standard null hypothesis testing framework, the two treatments had the statistically significant one-tailed probabilities of *p*(*C_i_*) = *p*(*E_j_*) = 0.95. In other words, the predictions of coexistence being made under an experimentally-parameterised BH model are not particularly strong, and, given the tenuous connection between predictions made in controlled pairwise experiments and observed patterns of co-occurrence in nature^7, 10, 11^, practitioners should exercise caution when using the approach employed herein to forecast the effects of climate change on communities.

Looking forward, researchers are encouraged to move from binary, all-or-nothing predictions of species coexistence to probabilistic, error-inclusive metrics more transparent in their predictions^12, 13^. Likewise, moving beyond phenomenological competition models of species interactions to more mechanistic formulations^14, 15^ will reduce the need for bias-prone model selection and permit an explicit accounting of the various limiting factors that give rise to niche and fitness differences between competitors.

## Acknowledgements

I thank Mary Van Dyke, Christopher Terry, Jürg Spaak, and Samuel Ross for constructive feedback and discussion.

## Author contributions statement

DWA performed all writing and analyses.

## Code availability

Code to replicate this analysis is available at https://doi.org/10.5281/zenodo.7460881

## Extended Data

**Fig. E1.**
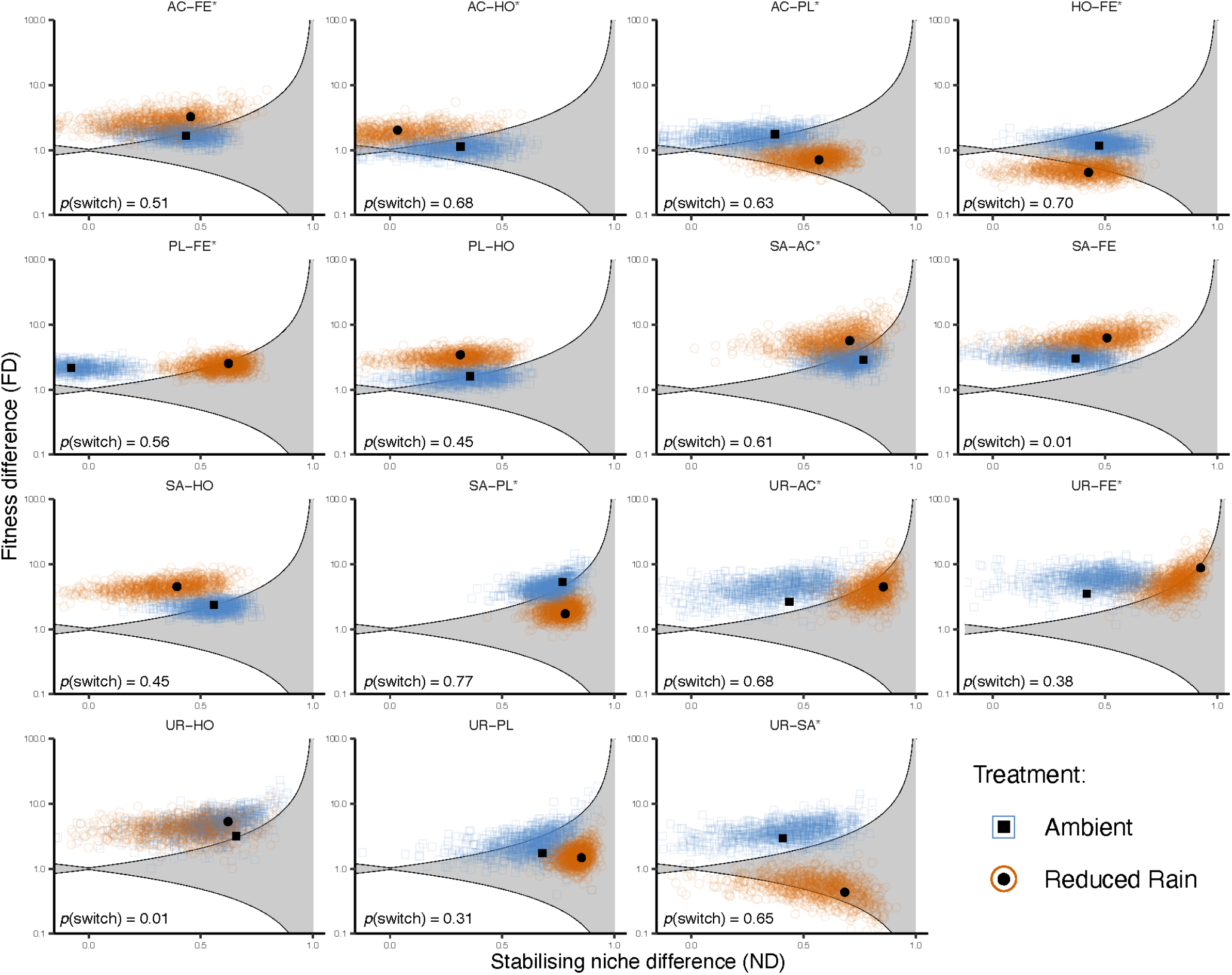
Predictions of coexistence outcomes for 15 species pairs. Points falling inside of the shaded region are those that satisfy the coexistence inequality. Colored points represent individual posterior draws from the authors’ original BH model (model 5). Black shapes show estimates from the BH model from the source publication. For each panel, the probability that a switch between coexistence and exclusion having occurred is also shown. Asterisks denote species pairs predicted to have experienced coexistence shifts in the original analysis.

